# Indian genetic heritage in Southeast Asian populations

**DOI:** 10.1101/2021.01.21.427591

**Authors:** Piya Changmai, Kitipong Jaisamut, Jatupol Kampuansai, Wibhu Kutanan, N. Ezgi Altınışık, Olga Flegontova, Angkhana Inta, Eren Yüncü, Worrawit Boonthai, Horolma Pamjav, David Reich, Pavel Flegontov

## Abstract

The great ethnolinguistic diversity found today in mainland Southeast Asia (MSEA) reflects multiple migration waves of people in the past. Deeply divergent East Eurasian hunter-gatherers were the first anatomically modern human population known to migrate to the region. Agriculturalists from South China migrated to the region and admixed with the local hunter-gatherers during the Neolithic period. During the Bronze and Iron Ages, the genetic makeup of people in MSEA changed again, indicating an additional influx of populations from South China. Maritime trading between MSEA and India was established at the latest 300 BCE, and the formation of early states in Southeast Asia during the first millennium CE was strongly influenced by Indian culture, and this cultural influence is still prominent today. Several ancient Indian-influenced states were located in present-day Thailand, and various populations in the country are likely to be descendants of people from those states. To systematically explore Indian genetic heritage in MSEA, we generated genome-wide SNP data (the HumanOrigins array) for 119 present-day individuals belonging to 10 ethnic groups from Thailand and co-analyzed them with published data from MSEA using the PCA, ADMIXTURE, *f_3_*-statistics, qpAdm, and qpGraph methods. We found South Asian low-level admixture in various MSEA populations which are probably descendants of people from the ancient Indian-influenced states, but failed to find a South Asian genetic component in present-day hunter-gatherer groups and relatively isolated groups from highlands in Northern Thailand. Our results also support close genetic affinity between Kra-Dai-speaking (also known as Tai-Kadai) and Austronesian-speaking populations, which fits a linguistic hypothesis suggesting cladality of the two language families.

**Author Summary:** Mainland Southeast Asia is a region with great ethnolinguistic diversity and complex population history. We studied genetic population history of present-day mainland Southeast Asian populations using genome-wide SNP data (the HumanOrigins array). We generated new data for 10 present-day ethnic groups from Thailand, which we further combined with published data from mainland and island Southeast Asians and worldwide populations. We revealed South Asian genetic admixture in various mainland Southeast Asian ethnic groups which are highly influenced by Indian culture, but failed to find it in groups who remained culturally isolated until recently. Our finding suggests that a massive migration of Indian people in the past was responsible for the spread of Indian culture in mainland Southeast Asia. We also found support for a close genetic affinity between Kra-Dai- and Austronesianspeaking populations, which fits a linguistic hypothesis suggesting cladality of the two language families.

## Introduction

Mainland Southeast Asia (MSEA) is a region with high ethnolinguistic diversity and complex population history. Hundreds of indigenous languages belonging to five language families (Austroasiatic, Austronesian, Hmong-Mien, Kra-Dai, and Sino-Tibetan) are spoken in MSEA [1]. Anatomically modern humans migrated to MSEA around 50000 – 60000 years ago [2]. Previous archaeogenetic studies indicate that the earliest MSEA individuals with genome-wide data available belong to the deeply diverged East Eurasian hunter-gatherers [3]. Andamanese hunter-gatherers (Onge and Jarawa) and MSEA Negritos are present-day populations related to the deeply diverged East Eurasian hunter-gatherers [3–4]. Neolithic populations in MSEA were established by admixture between these local hunter-gatherers and agriculturalists who migrated from South China around 4000 years ago [3–4]. The genetic makeup of MSEA Neolithic individuals is similar to present-day Austroasiatic-speaking populations [3–4]. That pair of seminal studies also detected additional waves of migrations from South China to MSEA during the Bronze and Iron Ages. Early states in MSEA during the first millennium CE, such as Pyu city-states, Funan, Dvaravati, Langkasuka, and Champa were established with a substantial influence from Indian culture [5]. There was evidence of Indian trading in MSEA and of glass bead manufacturing by MSEA locals using Indian techniques during the Iron Age [2]. Thailand is a country in the middle of MSEA, and many ancient Indianized states were located in its territory [5]. In Thailand 51 indigenous languages from five language families are attested.

We generated genome-wide SNP genotyping data for ten populations from Thailand: six Austroasiatic-speaking populations (Khmer, Kuy, Lawa, Maniq, Mon, and Nyahkur), one Hmong-Mien-speaking population (Hmong), one Kra-Dai-speaking population (Tai Lue), and two Sino-Tibetan-speaking populations (Akha and Sgaw Karen). Maniq, a MSEA Negrito group, are present-day hunter-gatherers. We combined our data with published MSEA and world-wide data. Our results revealed South Asian admixture in MSEA populations which are heavily influenced by Indian culture or which can be traced back to ancient Indianized states, and we failed to detect South Asian admixture in relatively isolated “hill tribes” (a term commonly used in Thailand for minority ethnic groups residing mainly in the northern and western highland region of the country) or in hunter-gatherers. The ubiquitous South Asian admixture in MSEA populations suggests a massive migration of South Asian populations to MSEA, which correlates with the spread of Indian culture across MSEA in the past. We also found Atayal-related ancestry in most Kra-Dai-speaking population in MSEA and South China, and that ancestry is absent in other MSEA groups apart from those with a clear history of Austronesian influence. The results suggest a link between Kra-Dai and Austronesian populations as previously suggested by linguistic studies proposing the existence of the Austro-Tai language macrofamily [6].

## Results

### Overview of the genetic make-up of ESEA populations

Using the HumanOrigins SNP array [7], we generated genome-wide genotyping data (~574,131 autosomal sites) for 10 present-day human populations from Thailand (Fig 1). We merged our data with published data for ancient and present-day worldwide populations (S1 table). To get an overview of population structure, we performed principal component analysis (PCA) (Fig 2). South Asian (SAS) populations lie on a well-known North-South cline [8]. Central Asian and Siberian populations lie between the European (EUR) – SAS and East-Southeast Asian (ESEA) clines. Mainland SEA Negritos occupied the space between the ESEA cline and the Andamanese (Onge). Munda populations, Austro-Asiatic-speaking populations from India which were shown in a previous study [9] to be a genetic mixture of South Asian and Southeast Asian populations, lie between the SAS and ESEA clines, as expected (Fig 2). Populations from East and Southeast Asia form a well-defined cluster, but positions of some populations such as Sherpa, Burmese, Mon, Thai, Cambodian, Cham, Ede, Malay, Khmer, Nyahkur, and Kuy are shifted towards the SAS cline (Fig 2).

**Fig 1.**
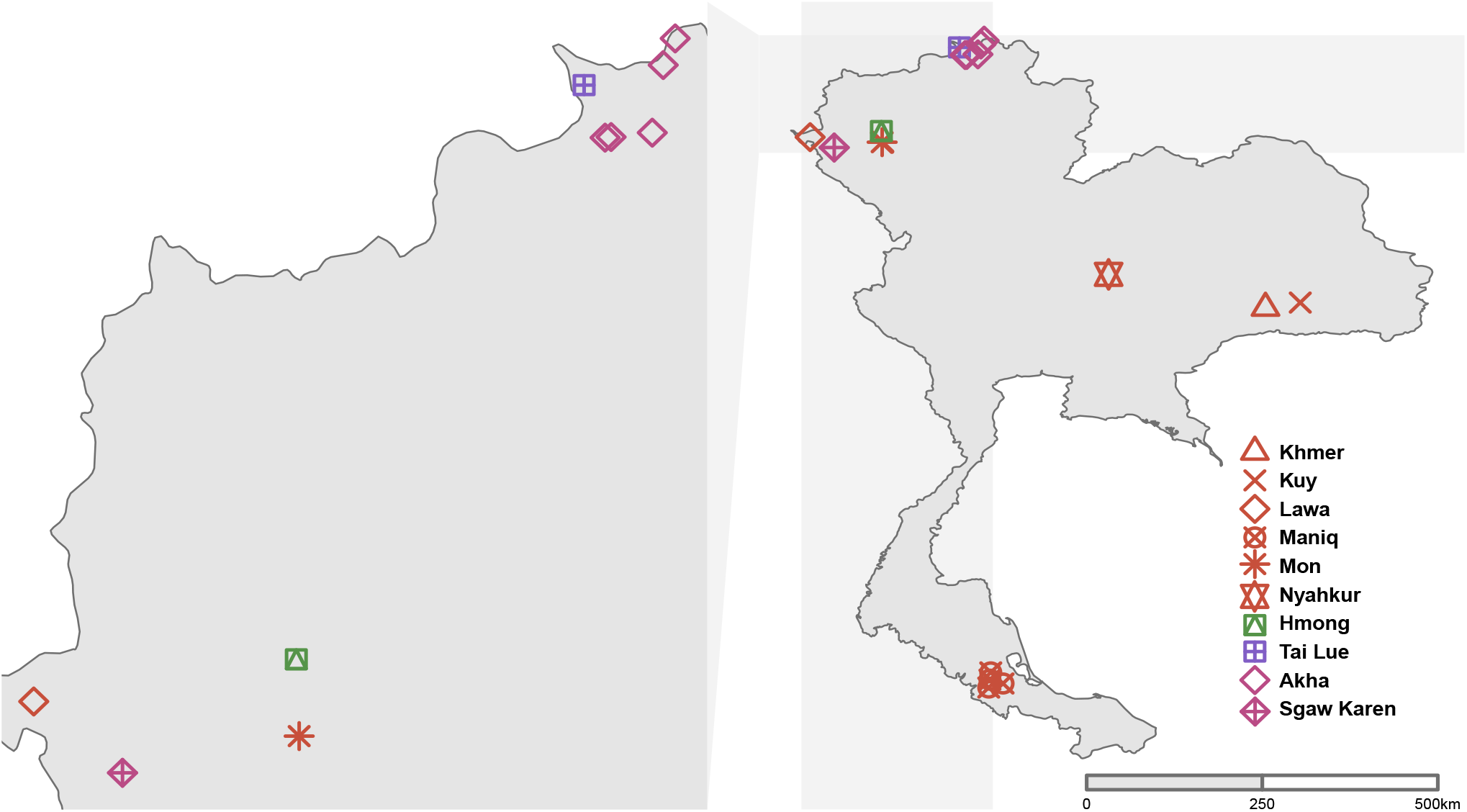
Locations of populations for whom genome-wide data was generated in this study. Colors represent language families: pink, Sino-Tibetan; green, Hmong-Mien; red, Austroasiatic; and purple, Kra-Dai. The map was created using R package “rwolrdmap” (https://cran.r-project.org/web/packages/rworldmap/).

**Fig 2.**
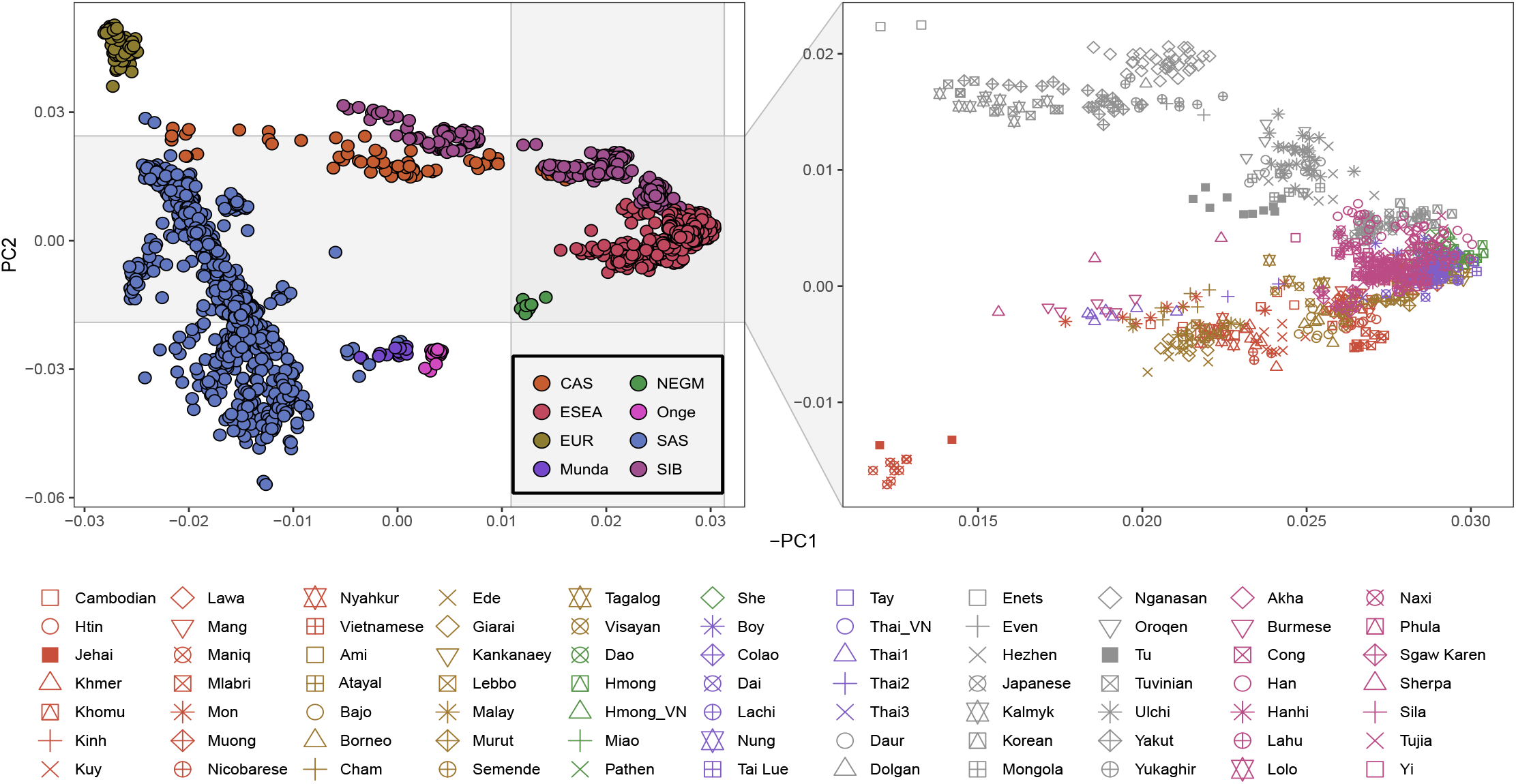
A principal component analysis (PCA) plot of present-day Eurasian populations. PCA was performed using PLINK. **Left panel**: An overview of the PC1 vs. PC2 space for all populations. The legend at the bottom of the plot lists abbreviations of meta-populations: CAS, Central Asians; ESEA, East and Southeast Asians; NEGM = Mainland Negritos; SAS, South Asians; EUR, Europeans; Munda, Austroasiatic-speaking populations (the Munda branch) from India; Onge, Onge (Andamanese hunter-gatherers); and SIB, Siberians. **Right panel**: A zoomed in view on the rectangle in the left panel.

Next, we performed a model-based clustering analysis using the ADMIXTURE approach. At 12 hypothetical ancestral populations, Burmese, Mon, Thai, Cambodian, Cham, Ede, Malay, Khmer, Kuy, and Nyahkur demonstrated an ancestry component (5% on average) which peaked in Indian populations (Fig 3). Due to the diversity of Thai (from Thailand) according to the PCA and ADMIXTURE analyses, we separated Thai into three groups labelled Thai1, Thai2, and Thai3. Their average SAS ancestry component proportions according to the ADMIXTURE analysis were as follows: 15, 7, and 3%, respectively (Figs 2 and 3).

**Fig 3.**
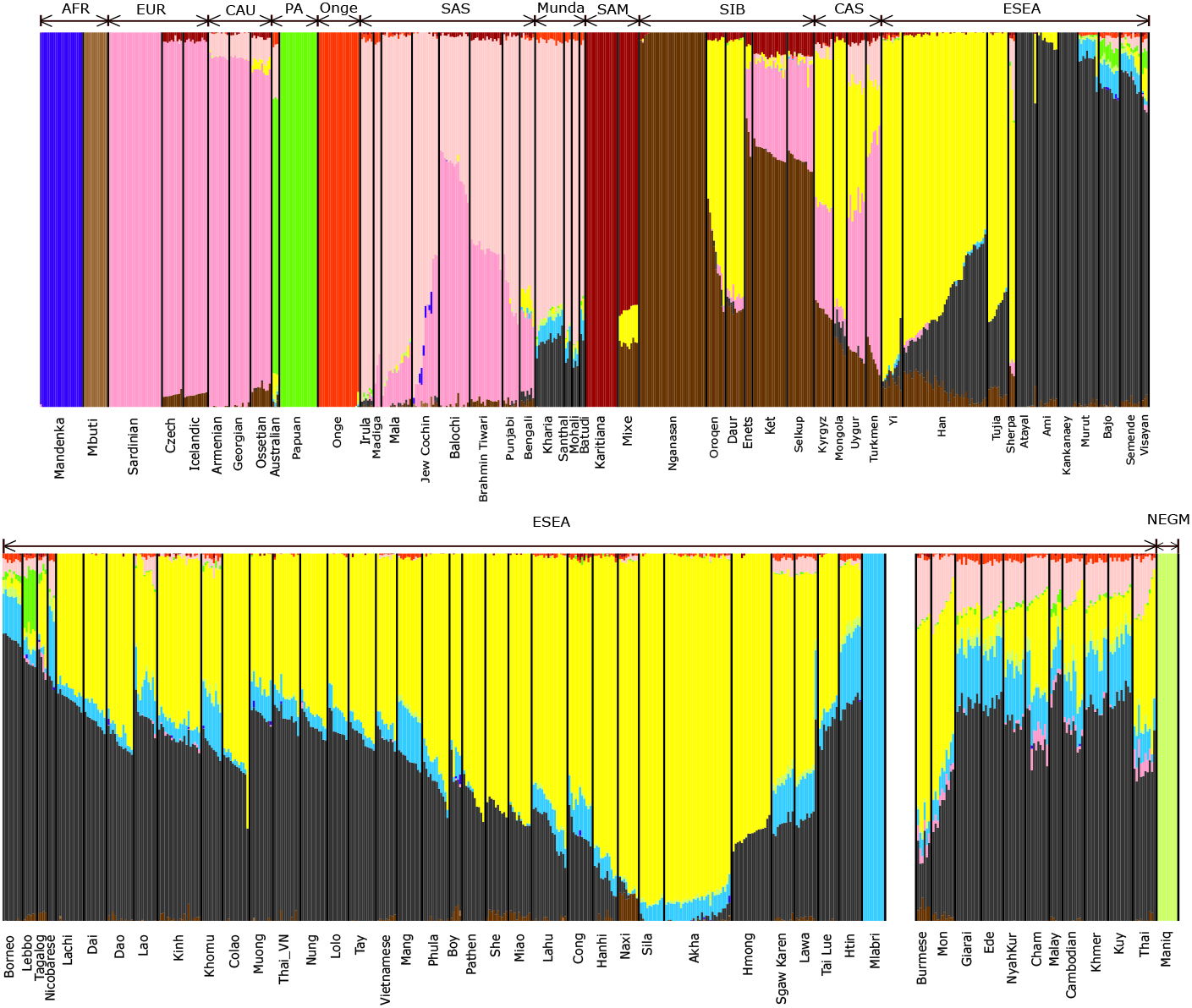
Results of an ADMIXTURE analysis. The plot represents results for 12 hypothetical ancestral populations. Abbreviations of meta-populations are shown above the plot: AFR, Africans; EUR, Europeans; CAU, Caucasians; PA, Papuans and Australians; Onge, Onge (Andamanese hunter-gatherers), SAS, South Asians; Munda, Austroasiatic-speaking populations (the Munda branch) from India; SAM, Native South Americans; SIB, Siberian; CAS, Central Asians; ESEA, East and Southeast Asians; and NEGM, Mainland Negritos.

Outgroup *f*_3_-statistics are used for measuring shared genetic drift between a pair of test populations relative to the outgroup population. We further explored hypothetical SAS admixture in MSEA by inspecting a biplot of outgroup *f*_3_-tests (Fig 4 and S1 Fig). We used Mbuti as an outgroup. On the y-axis, statistics *f*_3_(Mbuti; A, test group) are shown, where *A* are East Asian surrogates (Han or Dai) and “test groups” are other ESEA populations. On the x-axis statistics *f*_3_(Mbuti; B, test group) are shown, where *B* are South Asian populations (Brahmin Tiwari or Coorghi). In the coordinates formed by statistics *f*_3_(Mbuti; Han, test group) and *f*_3_(Mbuti; Brahmin Tiwari, test group) (Fig 4), most ESEA populations demonstrate a linear relationship between the genetic drift shared with Han and the drift shared with Brahmin Tiwari. However, positions of Burmese, Thai, Mon, Cham, Nyah Kur, Cambodian, Khmer, Malay, Giarai, and Ede are shifted from that main ESEA trend line. This shift can be interpreted as an elevated shared drift between the SAS group and the test population, as compared to other ESEA populations. Similar results were generated when we replaced Han and Brahmin Tiwari with Dai and Coorghi, respectively (S1 Fig).

**Fig 4.**
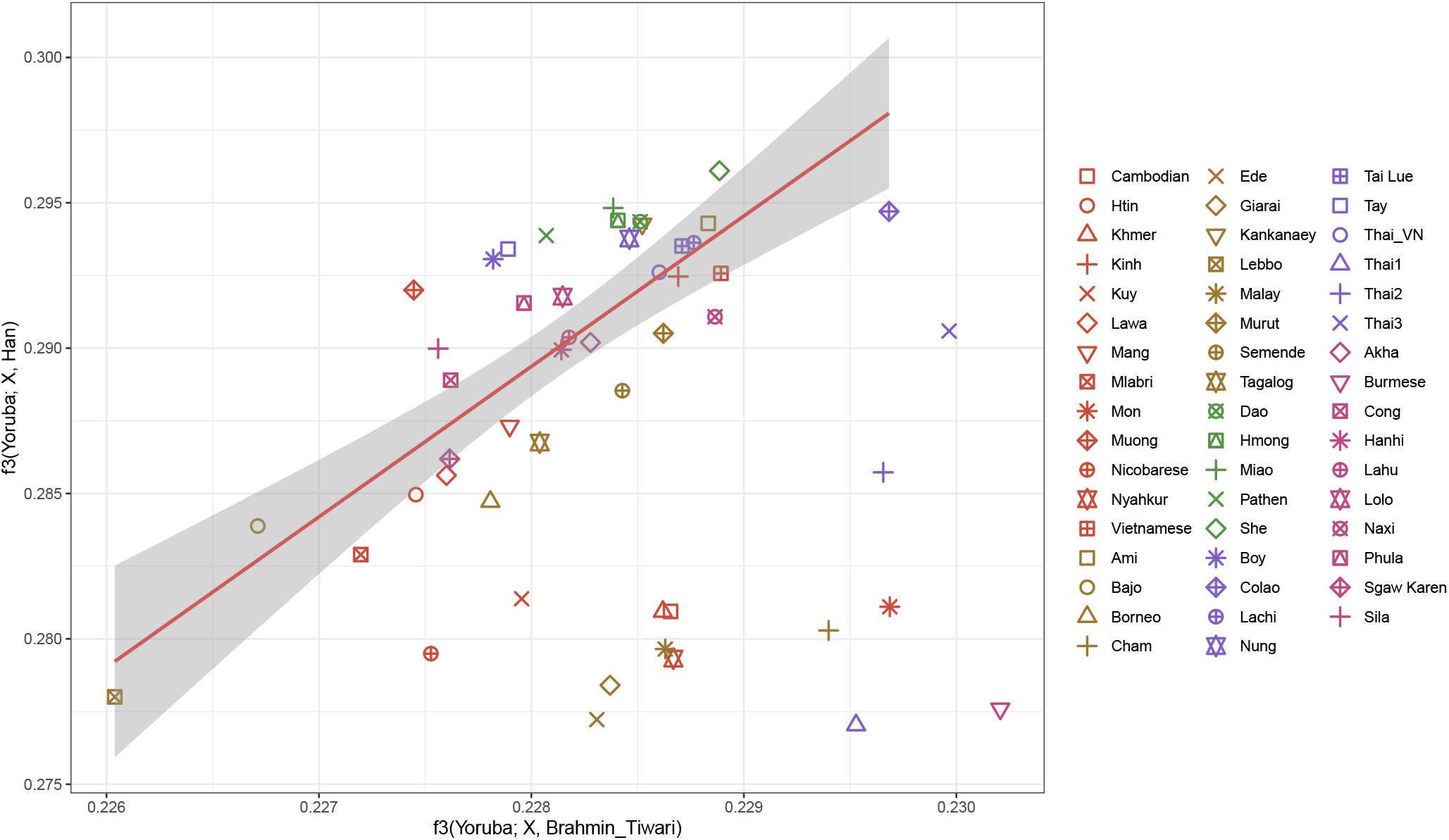
A biplot showing results of outgroup *f*_3_-tests. The biplot of *f*_3_(Mbuti; Brahmin Tiwari, X) vs. *f*_3_(Mbuti; Han, X) illustrates the amount of genetic drift shared between test ESEA populations and Brahmin Tiwari or Han. The trend line represents a ratio of shared genetic drifts that is common for most ESEA populations. The positions of few ESEA populations deviated from the trend line, which indicates elevated shared drift between the Indian reference population and the test population, as compared to most ESEA populations.

### Fitting admixture models using the qpWave, qpAdm, and qpGraph approaches

We also tested specific admixture models using the qpWave [10] and qpAdm methods [11, 12]. Previous studies indicate that deeply diverged East Eurasian hunter-gatherers (associated with the Hoabinhian archaeological culture), which are related to present-day Andamanese hunter-gatherers (Onge), were the first known anatomically modern humans who occupied MSEA [3, 4]. MSEA populations in the Neolithic period can be modelled as a mixture of local Hoabinhians and populations who migrated from East Asia [3, 4]. In our analysis, we used Atayal, Dai, and Lahu as ESEA surrogates. These populations speak languages which belong to three different language families: Austronesian, Kra-Dai, and Sino-Tibetan (the Tibeto-Burman branch), respectively. Onge was used as a surrogate for the deeply diverged East Eurasian hunter-gatherers. 55 populations composed of at least 5 individuals were used as South Asian surrogates. Outgroups (“right populations”) for all qpWave and qpAdm analyses were the following present-day populations: Mbuti (Africans), Palestinians, Iranians (Middle Easterners), Armenians (Caucasians), Papuans [7], Nganasans, Kets, Koryaks (Siberians), Karitiana (Native Americans), Irish, and Sardinians (Europeans).

We first explored cladality of population pairs using qpWave (Fig 5, S2 Table). In other words, we tested if one stream of ancestry from an ESEA surrogate is sufficient to model a Southeast Asian target population. We used a cut-off p-value of 0.05. We further tested 2-way and 3-way admixture models using qpAdm. We applied three criteria for defining plausible admixture models: a) all simpler models should be rejected according to the chosen p-value cutoff; b) the current model should not be rejected according to the chosen p-value cutoff; c) inferred admixture proportions ± 2 standard errors should lie between 0 and 1 for all ancestry components. If a model meets all three criteria, we consider the model as “fitting” or “passing” (S2 Table), although we caution that the only secure interpretation of qpWave or qpAdm tests is in terms of model rejection, and not model fit [13]. For testing 2-way and 3way admixture, we constructed models “ESEA + Onge” and “ESEA + Onge + South Asian”, respectively (Fig 5, S2 Table).

**Fig 5.**
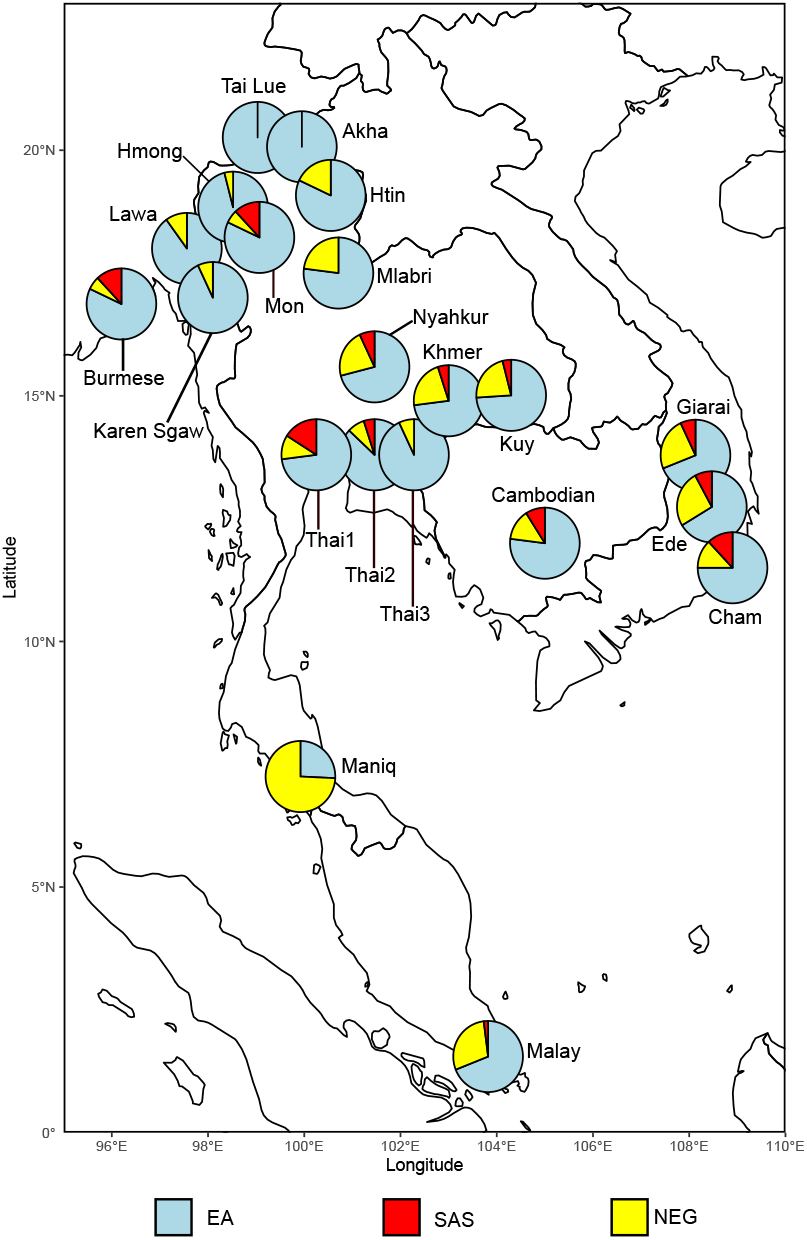
An overview of admixture proportions estimated by qpAdm. Admixture proportions were inferred using qpAdm with three groups of surrogates representing three ancestries: deeply diverged East Eurasian (NEG), South Asian (SAS), and East Asian (EA). Admixture proportions were averaged across all models which passed our criteria for “fitting” models. The map was plotted using R package “rnaturalearth” (https://github.com/ropensci/rnaturalearth).

Next, we tested more explicit demographic models using qpGraph. We first constructed two skeleton graphs using different SAS surrogates, Coorghi (Fig 6A) and Palliyar (Fig 6B). The worst residuals for the skeleton graphs were 2.43 and 2.24 SE intervals, respectively. Skeleton graph construction is explained in Materials and methods. We then exhaustively mapped target ESEA populations on all possible edges (except for edge0 in S2 Fig) on the skeleton graphs. We modeled the target populations as unadmixed (33 models per target population per skeleton graph), 2-way admixed (528 models per target population per skeleton graph), and 3-way admixed (5,456 models per target population per skeleton graph). We compared models with different numbers of admixture sources using a log-likelihood difference cut-off of 10 log-units or a worst residual difference cut-off of 0.5 SE intervals (see exploration of appropriate cut-offs on simulated genetic data in Ning et al., 2020 preprint [13]). For models with the same number of admixture sources, we used a log-likelihood difference cutoff of 3 log-units [14]. We also avoided models with trifurcations, i.e., when drift length on any “backbone” edge equals zero. Below we discuss best models found for the studied populations grouped by language family. The summary of qpWave, qpAdm, and qpGraph results is presented in Table 1. Full results are shown in S2 Table (qpWave and qpAdm) and S3 Table (qpGraph). S3 Table shows all qpGraph models satisfying the log-likelihood difference criteria. Edge number codes for the Coorghi and Palliyar skeleton graphs are illustrated in S2 Fig.

**Fig 6.**
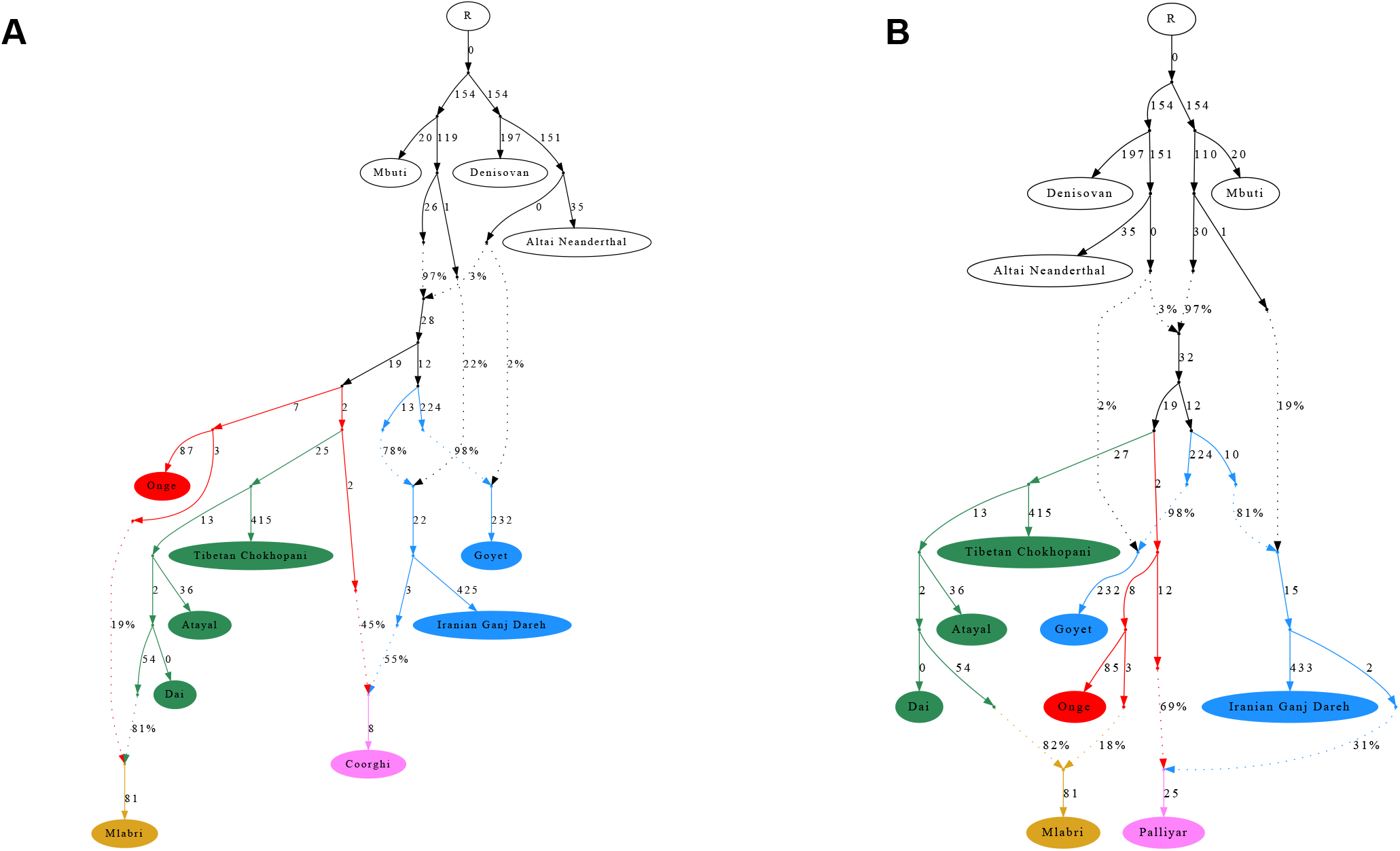
Skeleton graphs used for the qpGraph mapping method. We used the skeleton graphs to explore the genetic make-up of ESEA populations. We used different South Indian populations for two skeleton graphs: Coorghi in panel **A** and Palliyar in panel **B**.

**Table 1.** A summary of qpAdm and qpGraph admixture modelling results for the groups of interest. Labels of groups genotyped in this study are italicized.

| population | n | language family | country | qpAdm, best model | qpGraph, best model |
| --- | --- | --- | --- | --- | --- |
| Cambodian | 9 | Austroasiatic | Cambodia | ESEA + NEG + SAS | Atayal + Mlabri + SAS |
| Htin | 10 | Austroasiatic | Thailand | ESEA + NEG | Mlabri |
| *Khmer* | 10 | Austroasiatic | Thailand | ESEA + NEG or ESEA + NEG + SAS | Mlabri + SAS |
| *Kuy* | 10 | Austroasiatic | Thailand | ESEA + NEG or ESEA + NEG + SAS | Mlabri + SAS |
| *Lawa* | 10 | Austroasiatic | Thailand | ESEA + NEG | Tibetan + Mlabri |
| *Maniq* | 9 | Austroasiatic | Thailand | ESEA + NEG | Atayal + NEG |
| Mlabri | 10 | Austroasiatic | Thailand | ESEA + NEG | included in the skeleton graphs |
| *Mon* | 10 | Austroasiatic | Thailand | ESEA + NEG + SAS | before Tibetan + Mlabri (ESEA source) + SAS |
| *Nyahkur* | 9 | Austroasiatic | Thailand | ESEA + NEG + SAS | Mlabri + SAS |
| Cham | 10 | Austronesian | Vietnam | ESEA + NEG + SAS | Atayal + Mlabri + SAS (western source) |
| Ede | 9 | Austronesian | Vietnam | ESEA + NEG + SAS | Mlabri + SAS |
| Giarai | 11 | Austronesian | Vietnam | ESEA + NEG + SAS | Mlabri + SAS |
| Malay | 5 | Austronesian | Singapore | ESEA + NEG or ESEA + NEG + SAS | Atayal + Mlabri + SAS |
| *Hmong* | 10 | Hmong-Mien | Thailand | ESEA + NEG | before Atayal + Tibetan |
| *Tai Lue* | 9 | Kra-Dai | Thailand | ESEA | before Dai/Mlabri + Mlabri |
| Thai1 | 7 | Kra-Dai | Thailand | ESEA + NEG + SAS | before Tibetan + Mlabri (ESEA source) + SAS |
| Thai2 | 2 | Kra-Dai | Thailand | ESEA + NEG or ESEA + NEG + SAS | before Atayal + SAS |
| Thai3 | 1 | Kra-Dai | Thailand | ESEA or ESEA + NEG | Atayal + Mlabri |
| *Akha* | 31 | Sino-Tibetan | Thailand | ESEA | Tibetan + Mlabri (ESEA source) |
| Burmese | 6 | Sino-Tibetan | Myanmar | ESEA + NEG + SAS | Tibetan + Mlabri + SAS |
| *Sgaw Karen* | 10 | Sino-Tibetan | Thailand | ESEA + NEG | Tibetan + Mlabri |

#### Sino-Tibetan (Tibeto-Burman branch)

We studied Akha, Sgaw Karen from Thailand, and Burmese (15) from Myanmar. All three groups harbor ancestry from a Tibetan-related source (S3 Table). Akha was modeled as one stream of ancestry when Lahu was used as an ESEA surrogate in qpWave (S2 Table). Sgaw Karen requires an extra ancestry from the Onge surrogate in qpAdm analysis (S2 Table). The result agrees with qpGraph analysis where Sgaw Karen was modeled as a mixture of a Tibetan-related and a Mlabri-related source (S3 Table). Mlabri harbor a substantial proportion of deeply diverged East Eurasian ancestry (Fig 6). Additional gene flow from deep sources (edge7 and edge8) to Karen on the Coorghi skeleton decreased the worst residual by ~0.5 SE intervals, but the inferred admixture proportion was close to zero; therefore, these additional edges could be an artifact. Both qpAdm and qpGraph analyses indicated South Asian ancestry in Burmese: e.g. ~12% inferred by qpAdm (S2-3 Table). Burmese harbor ancestry from Tibetan-related + Mlabri-related + South Asian sources according to a best-fitting graph model (S3 Table).

#### Hmong-Mien

We analyzed Hmong from Thailand. We were not able to model Hmong as cladal with any of our three standard ESEA surrogates (Atayal, Dai, and Lahu). Then we tried to use Miao, a Hmong-Mien-speaking population from China, as an ESEA surrogate. We successfully modeled Hmong as Miao + Onge (S2 Table). The Hmong groups from Thailand and from Vietnam [16] are cladal according to qpWave (S2 Table). Our qpGraph result showed a low level of Tibetan-related ancestry (~2%) in Hmong (S3 Table).

#### Austronesian

There are four Austronesian-speaking populations included in this study: Cham, Ede (Rade), and Giarai (Jarai) from Vietnam [16], and Malay from Singapore [15]. qpAdm and qpGraph results revealed South Asian ancestry in all four Austronesian groups: 11.6%, 7.5%, 7.4%, and 2.1% in Cham, Ede, Giarai, and Malay, respectively, as inferred by qpAdm (S2-3 Table). Atayal is an Austronesian-speaking group from Taiwan, the homeland of Austronesian languages [17]. We failed to detect Atayal-related ancestry in Ede and Giarai S3 Table), while the ancestry is present in Cham and Malay. We found Mlabri-related ancestry in all four Austronesian-speaking populations (S3 Table)

#### Austroasiatic

We studied Htin [4], Khmer, Kuy, Lawa, Maniq, Mlabri [4], Mon, Maniq, and Nyahkur from Thailand, and Cambodians from Cambodia [7]. Maniq, a present-day hunter-gatherer Negrito group from Southern Thailand, has a major ancestry component derived from a deeply diverged East Eurasian group with ~74 % admixture proportion inferred by qpAdm (S2-3 Table). The ESEA source for Maniq is Atayal-related (S3 Table). Htin was modeled as a sister group of Mlabri by qpGraph (S3 Table). Both groups were modelled by qpAdm as having ESEA and Onge-related ancestry (S2-3 Table). Lawa was modeled as Mlabri-related + Tibetan-related ancestry (S3 Table). We detected South Asian admixture in five Austroasiatic-speaking groups in our study (Cambodian, Khmer, Kuy, Mon, and Nyahkur): 9.4%, 4.6%, 4.3%, 11.6%, and 7%, respectively, as inferred by qpAdm. Khmer, Kuy, and Nyahkur showed similar genetic makeups (S2-3 Table). We observed Atayal-related ancestry in Cambodian (S3 Table) and Tibetan-related ancestry in Mon, and these ancestry sources are rare in other Austroasiatic speaking populations.

#### Kra-Dai

We tested Kra-Dai-speaking populations from China (Dong, Dong Hunan, Gelao, Li, Maonan, Mulam, and Zhuang form Wang et al., 2020 preprint [18]) and Vietnam (Boy, Colao, Lachi, Nung, Tay, and Thai from Liu et al., 2020 [16]), and Thailand (Tai Lue from this study, Thai1, Thai2, and Thai3 from Lazaridis et al., 2014 [19]). Most of the Kra-Dai-speaking populations from China and Vietnam harbor Tibetan-related and Atayal-related ancestry (S3 Table). The Thai3 from Thailand was modelled as getting ~56% of its ancestry from a sister group of Atayal (S3 Table). Thai2 harbors ancestry from a source diverging before Atayal (S3 Table). Atayal-related ancestry is missing in Thai1 (S3 Table), but we found a source diverging before Tibetan Chokhopani when we mapped the Thai1 population on the Coorghi skeleton (S3 Table). We observed South Asian ancestry in Thai1 and Thai2, but that ancestry is missing in Tai Lue and Thai3 (S2-3 Table). qpAdm inferred South Asian admixture proportions in Thai1 and Thai2 at 17% and 5%, respectively.

## Discussion

Indian culture was long established in MSEA, which also influenced early states formation in the region during the first millennium CE [5]. Previous studies reported South Asian admixture in few populations from Southeast Asia [20–22]. Some studies used the same or similar populations as those in the current study but did not focus on South Asian admixture [16, 19, 23]. In this study, we thoroughly analyzed South Asian admixture in present-day Southeast Asia. We also investigated the genetic markup of populations in the region. Our results were consistent across various methods used in this study (ADMIXTURE, *f_3_*-statistics, qpAdm, qpGraph). There were just one or a few admixture graph models which fitted the data significantly better than ca. 6000 other models we tested per target population. qpAdm and qpGraph results were in agreement: adding a South Asian-related admixture edge never improved qpGraph model fits significantly when a 3-way model with South Asian admixture was rejected by qpAdm. We discuss the results by language family below.

### Sino-Tibetan (Tibeto-Burman branch)

Using qpWave, we were not able to reject cladality of Akha and Lahu, two Sino-Tibetan-speaking populations (S2 Table). Sgaw Karen required an extra stream of ancestry from an Onge-related population (S2 Table). The Onge-related ancestry in Sgaw Karen can be explained by admixture with an Austroasiatic-speaking population, which harbors high genetic ancestry from Hoabinhians[3, 4]. Our best-fitting admixture graph model for Sgaw Karen includes genetic contribution from a Mlabri-related group, which fits this explanation (S3 Table). The high worst residual of the best-fitting graph including Sgaw Karen probably reflects absence of an important ancestry source on our skeleton graph. Our qpAdm and qpGraph results consistently demonstrated that Burmese from Myanmar harbor ancestry from South Asian populations. All three Sino-Tibetan-speaking populations tested (Akha, Karen, and Burmese) have Tibetan ancestry according to the best-fitting qpGraph models (S3 Table).

### Hmong-Mien

The best-fitting qpAdm model for Hmong was Miao + Onge, with a minimal admixture proportion from the latter source. Cladality with Miao, another Hmong-Mien speaking population, was rejected (S2 Table). qpGraph modeling also indicated a low-level gene flow (~2%) from a sister group of Tibetan Chokhopani (S3 Table). The main ESEA ancestry for Hmong is a source diverging before Atayal (S3 Table).

### Austronesian

Malay from Singapore was modeled by qpGraph as a 3-way admixture involving sister groups of Atayal, Mlabri, and South Asian populations (S3 Table). Malay is an Austronesian language. It is not surprising that the Malay harbor some ancestry from a source related to Atayal, an Austronesian-speaking population from Taiwan. A previous study reported admixture from an Austroasiatic-speaking population in Austronesian populations from Indonesia [4]. We also detected the same signal in Malay, which is represented by ancestry from a sister group of Mlabri (S3 Table). Our results generated relying on various approaches indicate South Asian admixture in Malay and also in three other Austronesian-speaking populations from Vietnam, i.e., Cham, Ede, and Giarai (S2-3 Table). Y-haplogroups of West Eurasian origin (R1a-M420 and R2-M479) were reported in Ede and Giarai by Machold et al. 2019 [24], and Y-haplogroups R-M17 and R-M124 were reported in Cham by He et al. 2012 [25]. Using qpGraph, we were able to confirm the Atayal-related ancestry in Cham, but that gene flow signal was not supported in the case of Ede and Giarai (S3 Table). The results are consistent with a previous study by Liu et al. 2020 [16], which supports the spread of Austronesian language by cultural diffusion in Ede and Giarai.

### Austroasiatic

Htin can be modeled by qpGraph as a sister group of Mlabri (S3 Table). Both Mlabri and Htin languages belong to the Khmuic branch of the Austroasiatic family. A previous study showed that Mlabri has a genetic profile similar to early Neolithic individuals from mainland Southeast Asia [4]. The qpGraph best-fitting models for Maniq, a mainland Negrito group, incorporate 2-way admixture between an Atayal-related source and an Onge-related source, with predominant genetic contribution from the latter source. Even though Maniq speak an Austroasiatic language, a better surrogate for their ESEA source was Atayal, an Austronesian-speaking population (S3 Table). Maniq may harbor Atayal-related ancestry from Austronesian-speaking populations in Southern Thailand (where they reside) or from Malaysia nearby. Using qpGraph, we could model Lawa as a 2-way admixture between a sister group of Tibetan Chokhopani and Mlabri-related ancestry, with predominant contribution from the latter source (S3 Table). The Austroasiatic-speaking Lawa likely got Tibetan-related ancestry via Sgaw Karen. Around 1850, Sgaw Karen started migrating from present-day Myanmar to the region that was once exclusively occupied by Lawa [26]. There are villages where both Lawa and Sgaw Karen live alongside each other [27], and intermarriage between the two groups became more common recently [28]. A previous study [22] also observed genetic interaction between Karen and Lawa. We detected a minor South Asian admixture component (~4-5%) in Kuy using both qpAdm and qpGraph methods (S2-3 Table). Kutanan et al. 2019 [29] reported the presence of a West Eurasian Y-haplogroup R1a1a1b2a1b (R-Y6) in Kuy.

In this study, we generated new data for Austroasiatic-speaking Khmer from Thailand. Khmer is the official language of Cambodia, and Khmer is the majority population of Cambodia [1]. Our admixture graph modeling showed that Khmer from Thailand and Cambodians harbor two ancestry sources in common: a Mlabri-related source and South Asian ancestry (S3 Table). West Eurasian Y-haplogroups R1a1a1b2a2a (R-Z2123) and R1a1 were reported in Khmer [29] and Cambodians [30], respectively. The best-fitting model for Cambodians includes additional ancestry from an Atayal-related (i.e., Austronesian) source (S3 Table). Cambodians likely got this ancestry via Cham due to the long-lasting interaction between the ancient Cambodian and Cham Kingdoms [5]. Cham is also the largest ethnic minority in Cambodia today [1].

Mon and Nyahkur languages belong to the Monic branch of the Austroasiatic family [1]. Our qpGraph modeling found Mlabri-related and South Asian ancestry in both populations. A previous Y-chromosome study [29] reported various haplogroups of West Eurasian origin, such as J and R, in Mon, and haplogroup J2a1 (J-L26) in Nyahkur. The higher frequencies of West Eurasian Y-haplogroups in Mon correspond to the higher South Asian admixture proportion found in Mon as compared to Nyahkur. Mon harbors additional ancestry from a source close to Tibetan Chokhopani (S3 Table). Tibetan-related ancestry is missing in Nyahkur (S3 Table). The Nyahkur group is possibly a remnant of an ancient Monic-speaking population from the Dvaravati kingdom located within present-day Thailand [31]. Mon probably got the Tibetan-related ancestry via interactions with Sino-Tibetan-speaking populations in Myanmar. Most of present-day Mon in Thailand are descendants of refugees who migrated from Myanmar in the last few centuries [32]. There is some debate about the origin of Mon in the Lamphun province, whether they are the direct descendants of people from the ancient Mon state in present-day Thailand (ca. 1300 years before present), or their ancestors migrated from Myanmar in the last few hundred years. Our results favor the latter possibility due to the Tibetan-related genetic component found in Mon from Lamphun, which may reflect interaction with Burmese or other Sino-Tibetan-speaking populations in Myanmar where the density of Sino-Tibetan-speaking populations is much greater than in Thailand [1]. Furthermore, the Tibetan-related ancestry is absent in Nyahkur, another Monic-speaking population from Thailand.

### Kra-Dai

Atayal-related ancestry was found in most Kra-Dai-speaking populations in China and Vietnam, according to our analysis (S3 Table). We also observed Atayal-related ancestry in Thai3 from Thailand (S3 Table). Besides the Kra-Dai speakers, we were able to detect Atayal-related ancestry only in Austronesian-speaking populations (Malay, Cham) or non-Austronesian populations which have historical evidence of interactions with Austronesians such as Maniq and Cambodian (S3 Table). Furthermore, when we used Atayal as an ESEA surrogate in 3-way qpAdm models (ESEA + Onge + SAS), most of the models were rejected. Only the models with Thais (Thai1 and Thai2) as target populations were not rejected (S2 Table). The genetic link between Austronesian-speaking and Kra-Dai-speaking populations may reflect genealogical relationship of the two language families as suggested by the Austro-Tai hypothesis [6]. Tai Lue is one of Dai ethnic groups originating in South China [33]. The Tai Lue volunteers in our study migrated to Thailand less than a century ago from Myanmar. Cladality of Tai Lue with all three ESEA surrogates was not rejected using qpWave (S2 Table). However, qpGraph modeling supported a more complex model for Tai Lue: 2-way admixture between a source close to Dai and either a Mlabri-related or a source diverging before Atayal (S3 Table). The result suggests that after the migration from China, Tai Lue admixed with local MSEA populations, or that the genetic makeup of the Dai group that gave rise to the Tai Lue group studied here was different from the Dai groups sampled previously [34]. qpGraph modeling revealed different genetic makeups for the three Thai sub-groups delineated in this study (S3 Table). Both qpAdm and qpGraph methods consistently supported South Asian admixture in both Thai1 and Thai2 groups (S2-3 Table). Best-fitting models for Thai1 and Thai2 include different ESEA sources. The ESEA ancestry in the Thai1 group can be traced to a source close to Dai and possibly an additional source that diverged before Tibetans (S3 Table). The latter source may reflect admixture with a group that harbors a distinct ESEA source, such as Chinese Han. Chinese were estimated to comprise at least 10% of the Thailand population [35–36]. The Thai2 group was modelled having ESEA ancestry from a source close to Atayal (S3 Table). We failed to detect South Asian ancestry in Thai3, in contrast to Thai1 and Thai2. The best qpGraph model for the Thai3 group is a 2-way mixture between sister groups of Mlabri and Atayal (S3 Table). Our results revealed a considerable diversity of the Thai. Previous studies also reported differences in the genetic makeup of the Thai from different locations [20, 22, 29]. Samples of all Thai individuals included in this study were obtained from the European Collection of Cell Cultures, and we cannot trace the origin of the samples in that collection [19]. Systematic sample collection at various locations will likely provide insight into the genetic diversity of the Thai.

Our study revealed substantial South Asian admixture in various populations across Southeast Asia (~2-16% as inferred by qpAdm). We observed South Asian admixture in some populations (Cham, Ede, Giarai, Khmer, Kuy, Nyahkur, and Thai) for whom the admixture was not reported before [16, 19, 23]. Most populations harboring South Asian admixture were heavily influenced by Indian culture in the past or are related to descendants of ancient Indianized states in Southeast Asia. In contrast, we failed to detect South Asian admixture in most “hill tribes” and in present-day hunter-gatherer groups from Thailand. Consequently, the spread of Indian influence in the region can be explained by extensive movement of people from India rather than by cultural diffusion only. The distance from the coast may affect South Asian gene flow as central and southern Thai harbor South Asian ancestry, but the ancestry is missing in northern Thai, who reside a long distance from the sea [22]. In this study, we also observed genetic diversity in Thai, but the exact location of the Thai individuals analyzed here is unknown. We detected subtle differences in populations with similar ethnolinguistic backgrounds, such as Khmer from Thailand and a Khmer-speaking population (Cambodian) from Cambodia. We observed Atayal-related ancestry (~3-38% as inferred by qpGraph) in most Kra-Dai-speaking populations from China, Vietnam and in one group from Thailand. The results suggest a genetic connection between Austronesian and Kra-Dai-speaking populations.

## Materials and methods

### Sampling

Sample collection and DNA extraction for all new Thailand populations in this study apart from Akha was described in previous studies [23, 37–41]. Saliva samples were obtained from volunteers with signed informed consent from four Akha villages in the Chiang Rai province, Thailand. The study was approved by the Ethic Committee of Khon Kaen University. We performed DNA extraction as described elsewhere [42]. See a list of individuals for whom genetic data is reported in this study in S4 Table.

### Dataset preparation

Diploid genome-wide SNP data was generated using the HumanOrigins SNP array [7]. We merged the new data with published ancient and present-day world-wide populations (S1 Table) using PLINK v. 1.90b6.10 (https://www.cog-genomics.org/plink/). We first combined present-day populations and applied a per site missing data threshold of 5% to create a dataset of 574,131 autosomal SNPs. We then added data from ancient populations. The Upper Paleolithic individual from Goyet had the highest missing data percentage per individual (30%). We used the dataset for all analyses except for ADMIXTURE.

### PCA

The principal component analysis (PCA) was performed using PLINK v. 1.90b6.10 (https://www.cog-genomics.org/plink/) on selected populations (S1 Table) from the following regions: Central, East, Southeast, and South Asia, Andamanese Islands, Siberia, and Europe.

### ADMIXTURE

We performed LD filtering using PLINK v. 1.90b6.10 with the following settings: window size = 50 SNPs, window step = 5 SNPs, r^2^ threshold = 0.5 (the PLINK option “--indep-pairwise 50 5 0.5”). LD filtering produced a set of 270,700 unlinked SNPs. We carried out clustering analysis using ADMIXTURE v. 1.3 (https://dalexander.github.io/admixture/download.html), testing from 8 to 13 hypothetical ancestral populations (K) with tenfold cross-validation. We performed five iterations for each value of K. We selected K = 12 for presentation according to the highest model likelihood. We further ran up to 30 iterations for K = 12 and ranked them by model likelihood.

### Outgroup *f*_3_-statistics

We computed *f*_3_-statistics [7] using qp3Pop v. 420, a software from the ADMIXTOOLS package (https://github.com/DReichLab/AdmixTools). We ran *f*_3_(Mbuti; X, test group), where X are East Asian surrogates (Han or Dai) or South Asian (Brahmin Tiwari or Coorghi) surrogates. The test groups are various ESEA populations.

### qpWave and qpAdm

We used qpWave v. 410 and qpAdm v. 810 from the ADMIXTOOLS package. We used the following populations as outgroups (“right populations”) for all qpWave and qpAdm analyses: Mbuti (Africans), Palestinians, Iranians (Middle Easterners), Armenians (Caucasians), Papuans [7], Nganasan, Kets, Koryaks (Siberians), Karitiana (Native Americans), Irish, and Sardinians (Europeans). We used Atayal, Dai, and Lahu as ESEA surrogates. We used Onge as a surrogate for the deeply diverged East Eurasian hunter-gatherers. We used 55 different populations as alternative South Asian surrogates (S2 Table).

We tested a pair of a test population and an ESEA surrogate using qpWave. We used a cut-off p-value of 0.05 for qpWave modeling. We performed 2-way and 3-way admixture modeling using qpAdm. 2-way admixture was modeled as “target population = ESEA surrogate + Onge”, and 3-way admixture was modeled as “target population = ESEA surrogate + Onge + SAS surrogate”. We applied three criteria for defining plausible admixture models: a) all simpler models should be rejected according to the chosen p-value cutoff; b) the current model should not be rejected according to the chosen p-value cutoff; c) inferred admixture proportions ± 2 standard errors should lie between 0 and 1 for all ancestry components.

### qpGraph

We used qpGraph v. 6412 from the ADMIXTOOLS package with the following settings: outpop: NULL, blgsize: 0.05, lsqmode: NO, diag: 0.0001, hires: YES, initmix: 1000, precision: 0.0001, zthresh: 0, terse: NO, useallsnps: NO. We used the following criteria to select the best-fitting model. Models with different numbers of admixture sources were compared using a log-likelihood difference cut-off of 10 log-units or a worst residual difference cut-off of 0.5 SE intervals [13]. We used a loglikelihood difference cut-off of 3 log-units for models with the same number of parameters [14].

We started with the following five populations: Denisovan (archaic human), Altai Neanderthal (archaic human), Mbuti (African), Atayal (East Asian), and Goyet (ancient West European hunter-gatherer). A best-fitting model is illustrated in S3 Fig. We fixed Neanderthal-related (node nA in S3 Fig) admixture proportion in non-Africans at 3%. Goyet requires extra admixture from this Neanderthal-related source. When this admixture edge was missing, the worst *f*-statistic residual increased from 2.13 to 4.56. We further mapped additional populations on the graph, one at a time. We mapped a new population on all possible edges on the graph as unadmixed, 2-way, and 3-way admixed. We mapped Onge on the 5-population graph (S3 Fig) and then Dai on the 6-population skeleton graph (S4 Fig). Best-fitting graphs including Onge and Dai are shown in S4 Fig and S5 Fig, respectively.

We further mapped an ancient Iranian herder individual from Ganj Dareh (I1947 [8]). A best-fitting model for this individual is a 2-way mixture between a putative West Eurasian source and a basal Eurasian source (S6 Fig). Basal Eurasian admixture in ancient Iranians was reported in a previous study [43]. Mlabri can be modeled as ESEA + Onge-related sources (S7 Fig), which is consistent with a previous study [4].

We mapped South Asian populations, Coorghi or Palliyar, on the graph in S7 Fig. Both populations can be modeled as a 2-way mixture between ancient Iranian-related and deep-branching East Eurasian sources (S8A and B Fig). The positions of the deep East Eurasian source for Coorghi and Palliyar are slightly different, but both are among the deepest East Eurasian branches.

Next, we added an ancient individual, Tibetan Chokhopani from Nepal (S1 Table), as the last population on the skeleton graphs. The best-fitting model for this individual was an unadmixed branch in the ESEA clade before the divergence of Atayal (Fig 6A and B). The total numbers of SNPs used for fitting the skeleton graphs with Coorghi and Palliyar were 311,259 and 317,327, and the worst absolute *f*_3_-statistic residuals were are 2.43 and 2.24 SE, respectively.

We mapped present-day target populations on all possible edges (except for edge0 in S2 Fig) on the skeleton graphs as unadmixed, 2-way admixed, and 3-way admixed. In total, we tested 6,017 models per target population per skeleton graph.

## Data Availability

Genome-wide genotyping data generated for this study will be made publicly available when the manuscript is published.

## Acknowledgments

We thank all volunteers in Thailand who donated the samples for our study. We thank Phangard Neamrat, Jaeronchai Chuaychu, and Prateep Panyadee for assisting with sample collection. This work was supported by the Czech Ministry of Education, Youth and Sports: 1) Inter-Excellence program, project #LTAUSA18153; 2) Large Infrastructures for Research, Experimental Development and Innovations project “IT4Innovations National Supercomputing Center – LM2015070”. E.Y., O.F., P.C., and P.F., were also supported by the Institutional Development Program of the University of Ostrava. J.K. acknowledges partial support provided by Chiang Mai University, Thailand.

## Supporting information

**S1 Fig.**
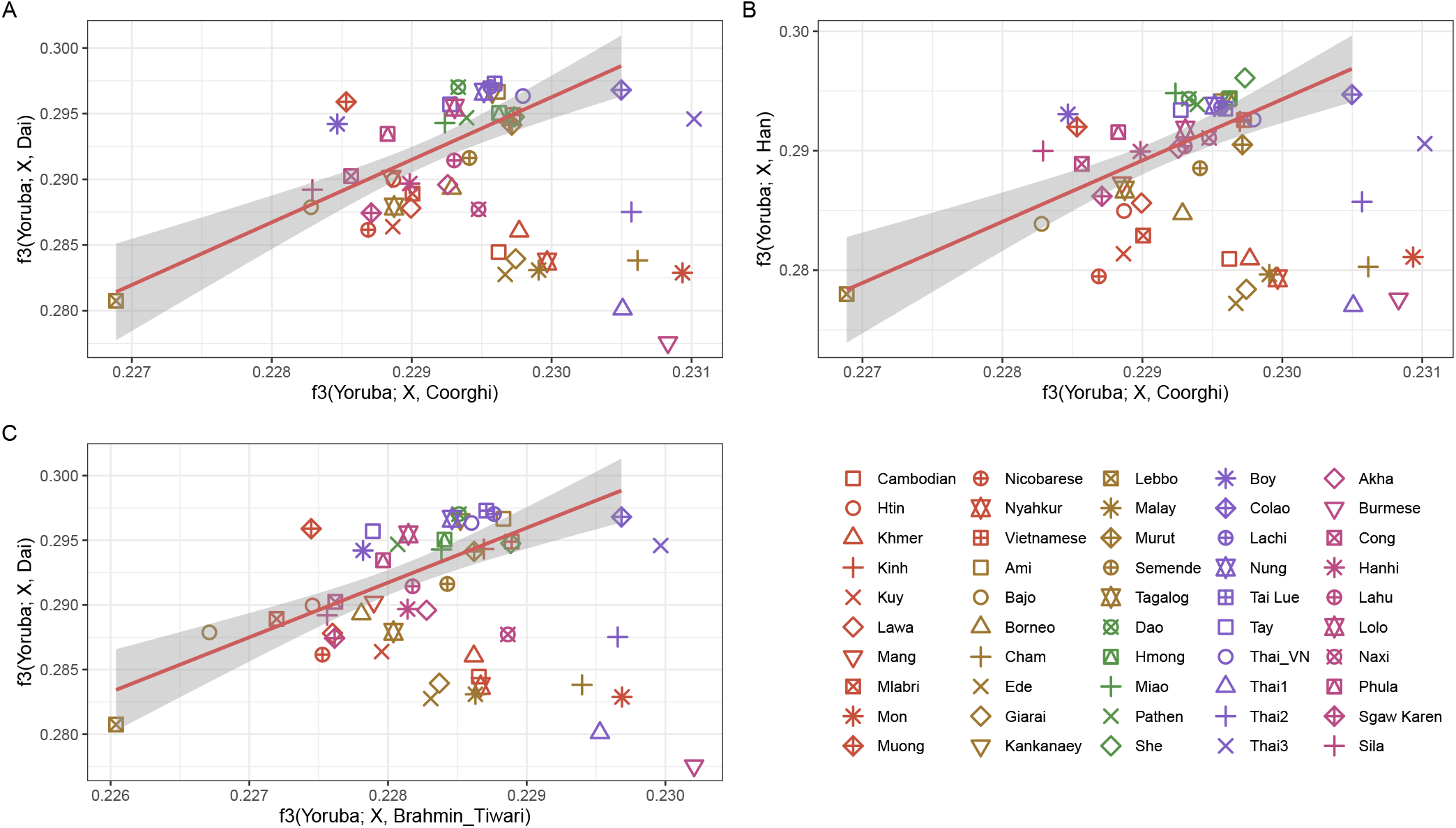
**A biplot of** *f_3_*(Mbuti; Coorghi, X) vs. *f_3_*(Mbuti; Dai, X) (A), *f_3_*(Mbuti; Coorghi, X) vs. *f_3_*(Mbuti; Han, X) **(B)**, and *f_3_*(Mbuti; Brahmin Tiwari, X) vs. *f_3_*(Mbuti; Dai, X) (**C**).

**S2 Fig.**
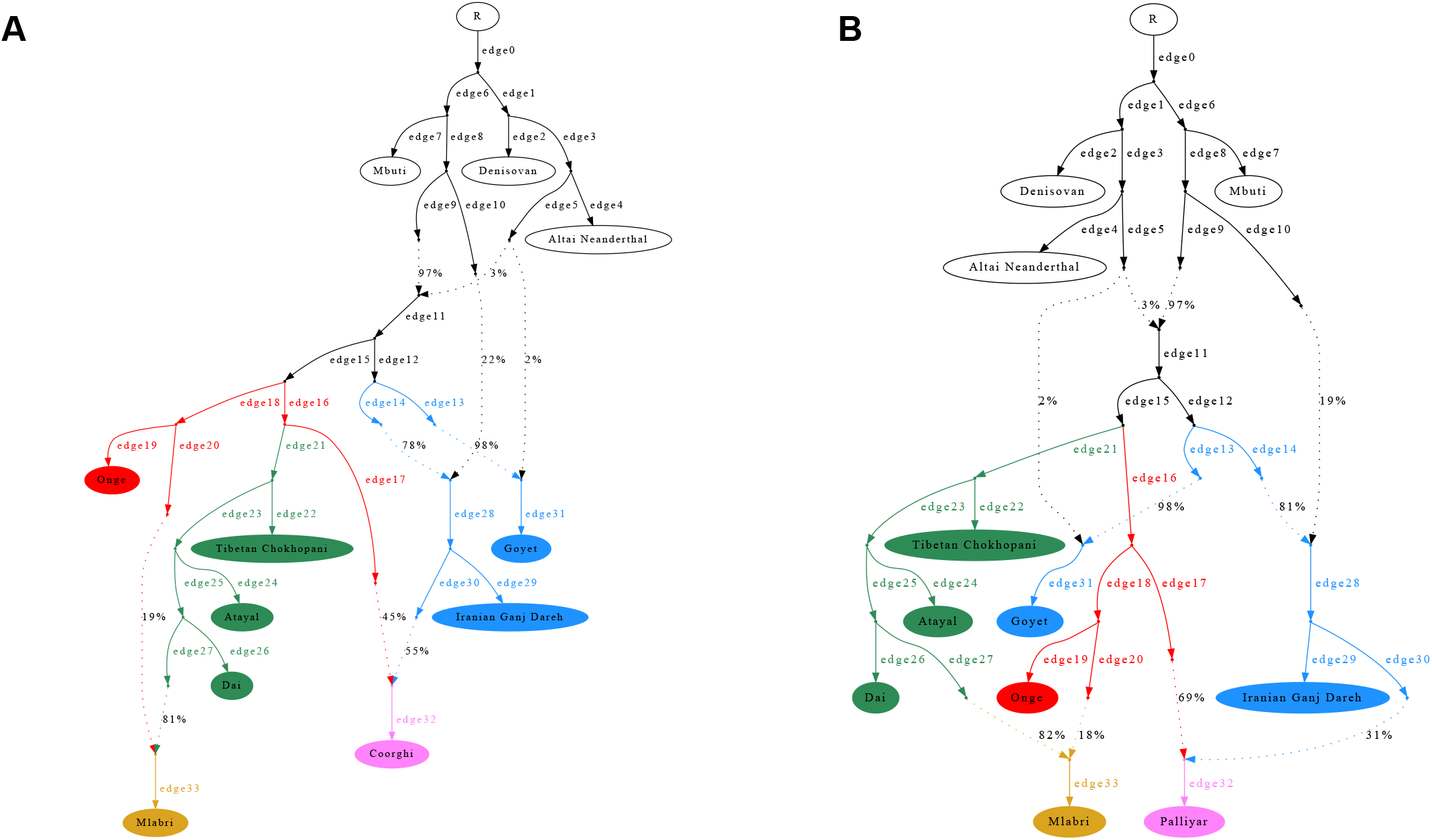
Skeleton graphs used for qpGraph mapping, with edges numbered. Coorghi was used as an Indian surrogate for skeleton graph **A** and Palliyar for skeleton graph **B**.

**S3 Fig.**
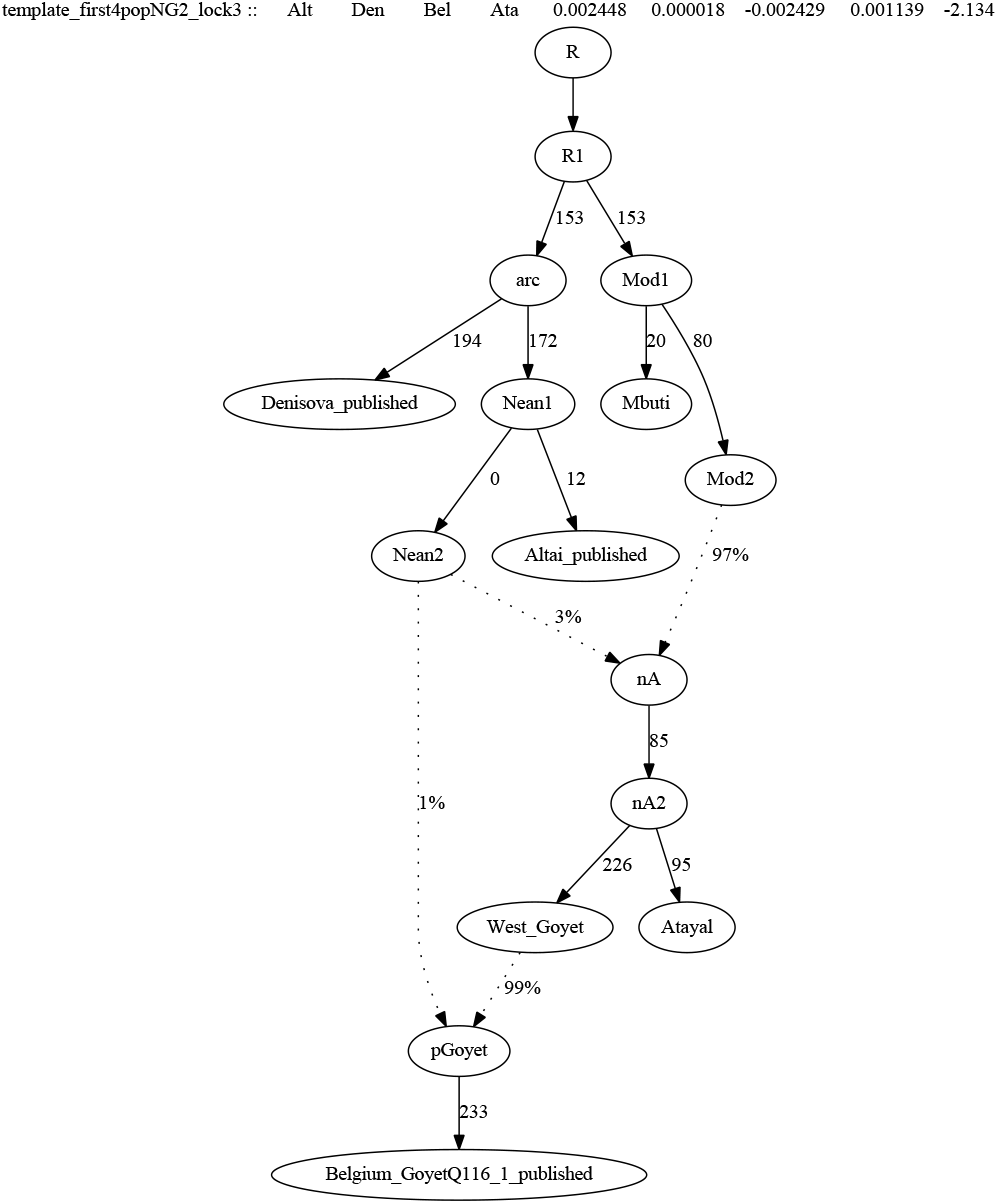
The starting skeleton graph with 5 populations.

**S4 Fig.**
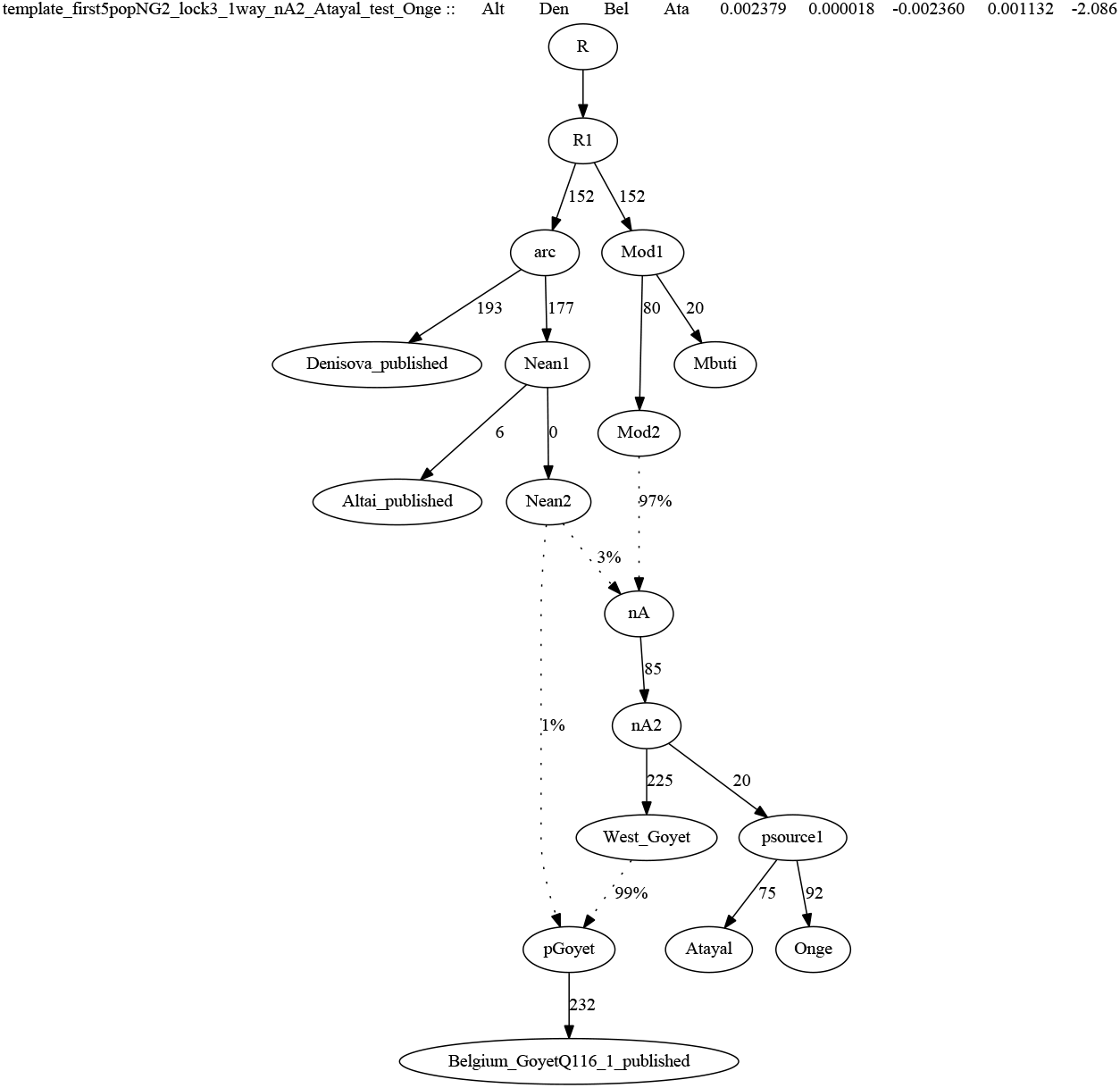
The best-fitting graph for Onge mapped on the 5-population skeleton graph (Fig S3).

**S5 Fig.**
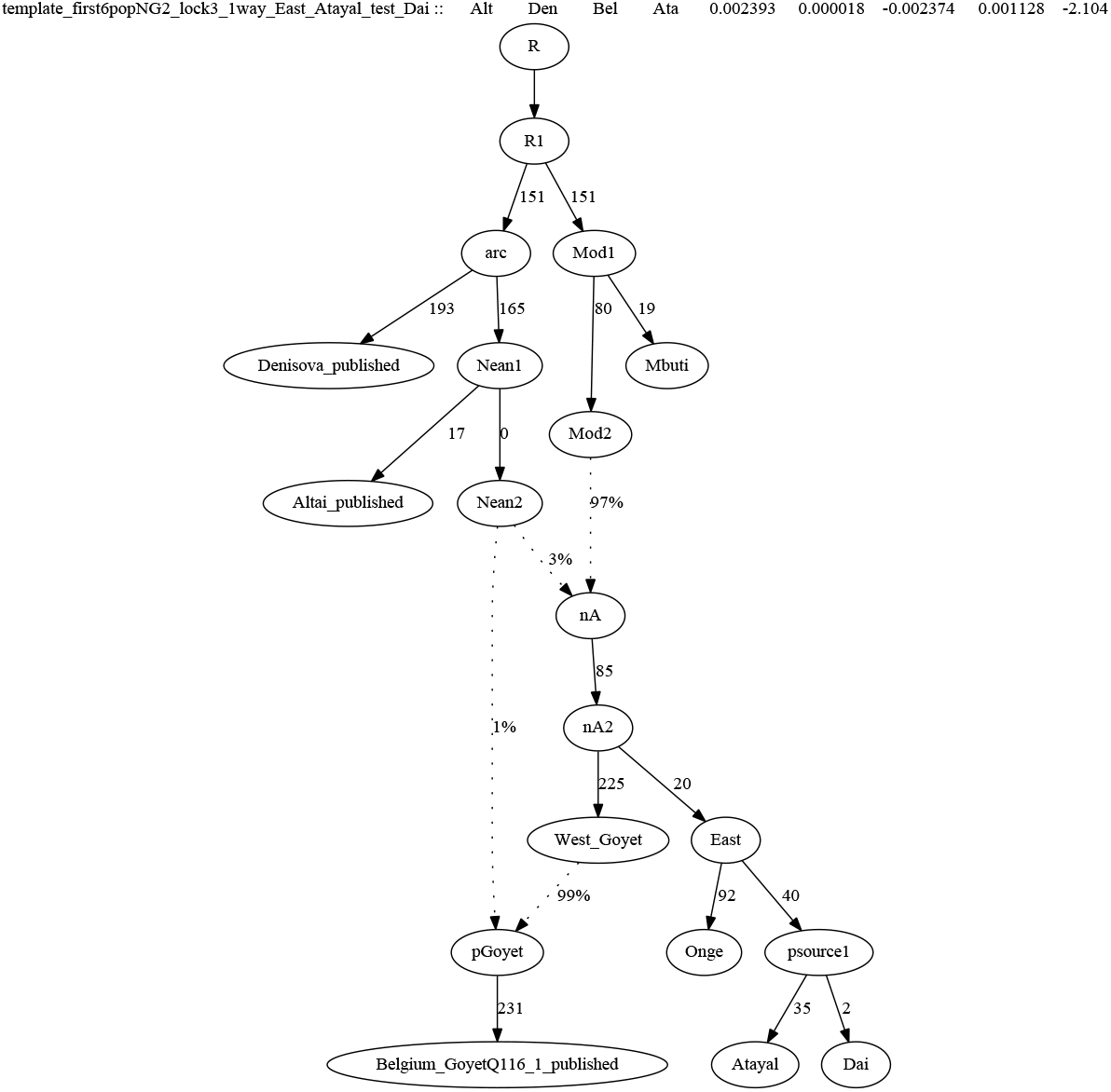
The best-fitting graph for Dai mapped on the 6-population skeleton graph (Fig S4).

**S1 Fig.**
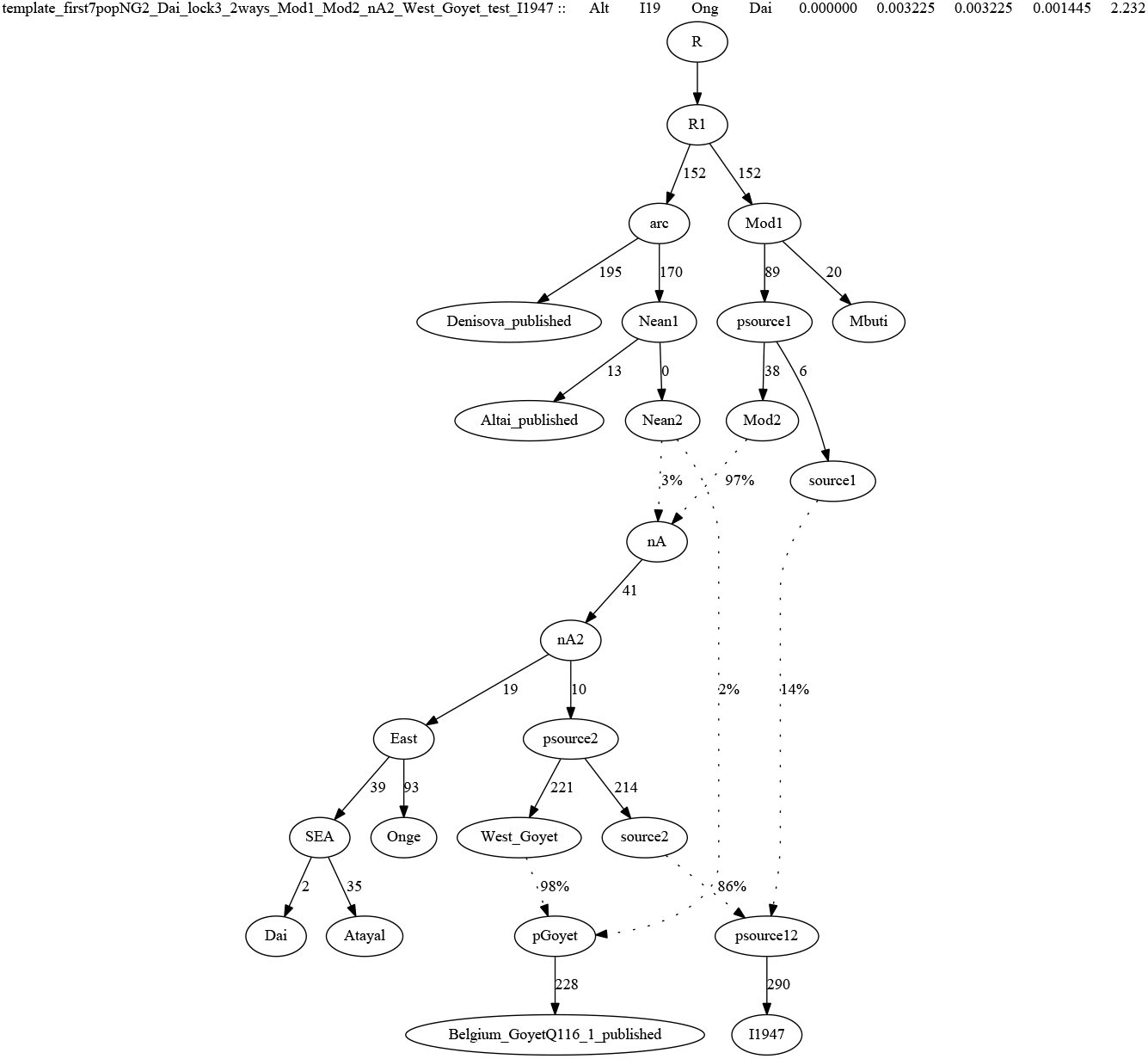
The best-fitting graph for an ancient Iranian herder form Ganj Dareh mapped on the 7-population skeleton graph (Fig S5).

**S7 Fig.**
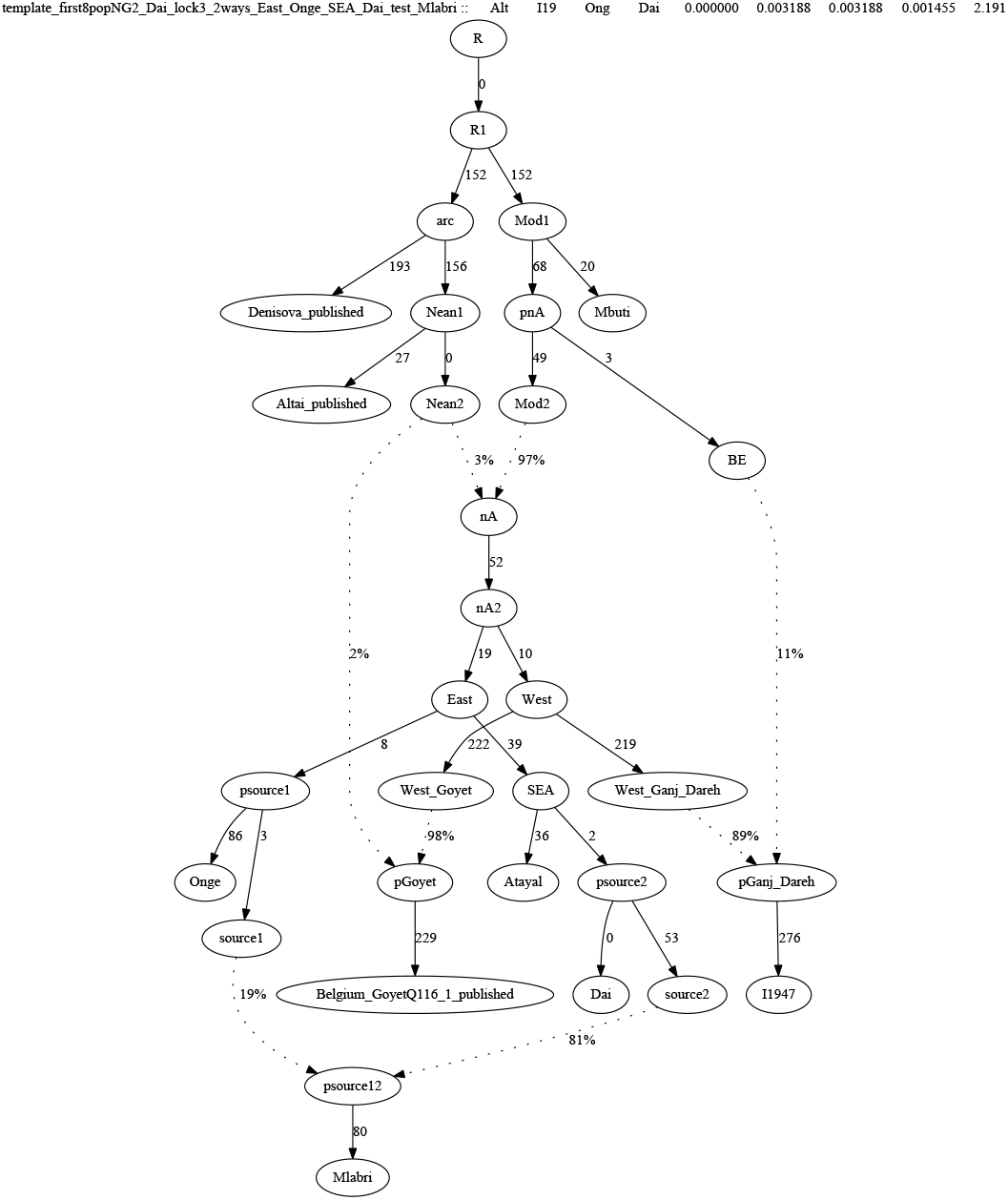
The best-fitting graph for Mlabri mapped on the 8-population skeleton graph (Fig S6).

**S8 Fig.**
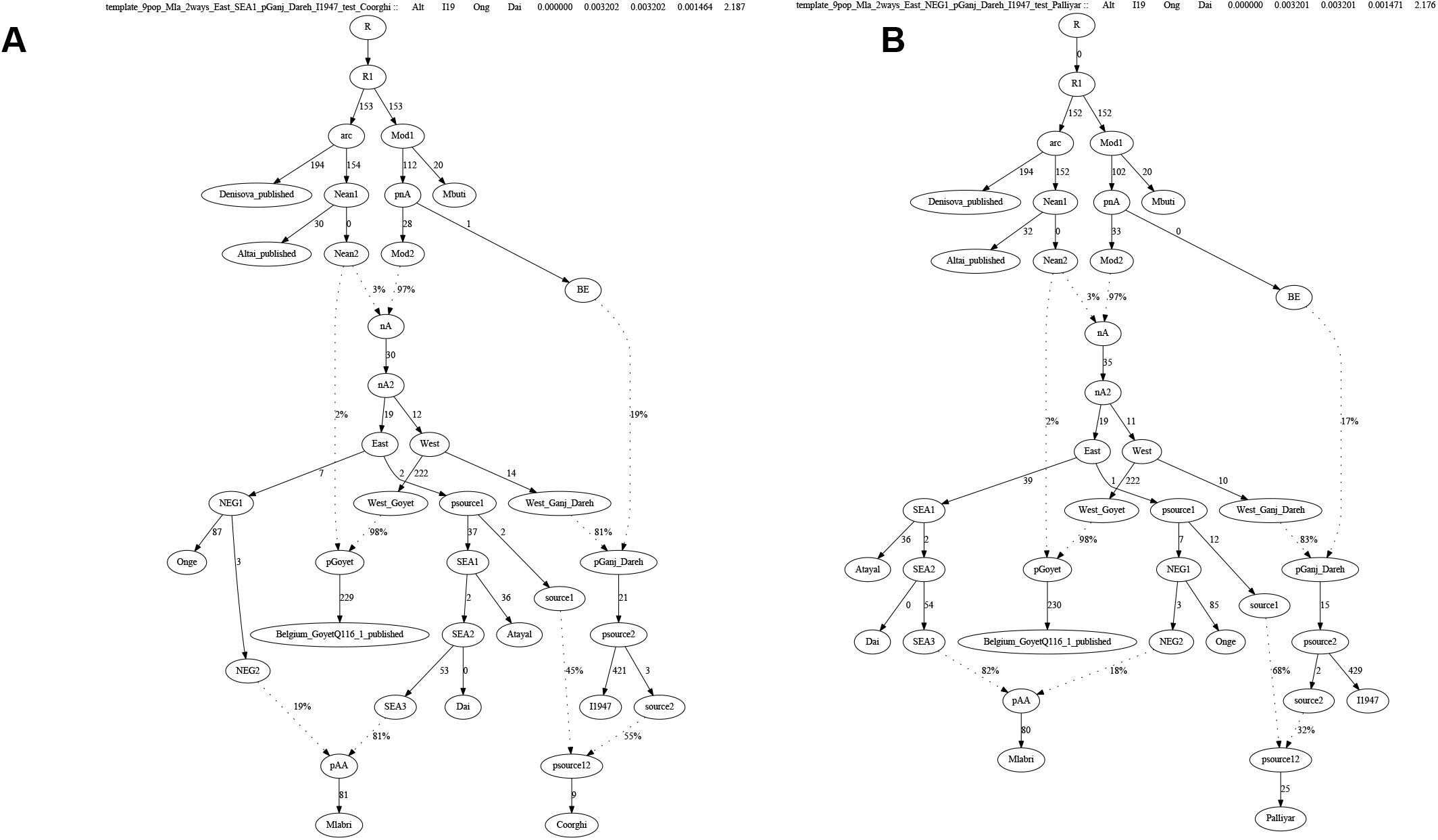
The best-fitting graphs for Coorghi (A) and Palliyar (B) mapped on the 9-population skeleton graph (Fig S7).

**S1 Table. Information on reference populations used in this study.**

**S2 Table. qpWave and qpAdm results.**

**S3 Table. All best-fitting qpGraph models.**

**S4 Table. Metadata for newly genotyped present-day individuals.**

